# Synaptic vesicle proteins are selectively delivered to axons in mammalian neurons

**DOI:** 10.1101/2022.02.08.479521

**Authors:** Emma T. Watson, Michaela M. Pauers, Jason D. Vevea, Edwin R. Chapman

## Abstract

Neurotransmitter-filled synaptic vesicles (SV) mediate synaptic transmission and are a hallmark specialization in neuronal axons. Yet, how SV proteins are sorted to presynaptic nerve terminals remains the subject of debate. The leading models posit that these proteins are randomly trafficked throughout neurons and are selectively retained in presynaptic boutons. Here, we used the RUSH system, in conjunction with HaloTag labeling approaches, to study the egress of two distinct SV proteins from the soma of cultured neurons. In sharp contrast to the selective retention model, both proteins selectively and specifically entered axons and did not traffic through dendrites; only upon overexpression do SV proteins spillover into other compartments. Moreover, we observed that SV constituents were first delivered to the presynaptic plasma membrane before incorporation into SVs. These experiments reveal a new-found membrane trafficking pathway in classically polarized mammalian neurons and provide a glimpse at the first steps of SV biogenesis.

## Introduction

Neurons present a dramatic example of cell polarization. These highly specialized and asymmetric cells form elaborate axonal and dendritic arbors, with some axons extending great distances (e.g., axons in a blue whale can reach a length of thirty meters (Smith 2009)). Within this polarized framework, axons and dendrites are highly adapted to carry out different functions and, consequently, each harbor somewhat distinct molecular constituents. For example, in chemical synapses, dendrites require a steady supply of receptors and proteins that are involved in post-synaptic signaling cascades, whereas axons require the machinery that drives the synaptic vesicle (SV) cycle, including the exocytosis of neurotransmitters. How this molecular and cellular polarity is maintained, specifically in the case of highly extended axons, is an essential question since preserving this extreme polarity underlies neuronal function. Indeed, defects in axonal transport have been implicated in a variety of neurodegenerative diseases (Hung and Link 2011; Maday et al. 2014; May-simera and Liu 2013; Vicario-Orri et al. 2014).

Many aspects of axonal and dendritic transport are well characterized (Hirokawa 1993; Maday et al. 2014; Subhojit Roy 2014; Twelvetrees 2020). Specifically, several families of motor proteins, which carry transport vesicles along microtubule and actin tracks, have been described (Nobutaka et al. 2005; Kneussel and Wagner 2013). Together, motor proteins and the cytoskeleton constitute a transport network that supports the formation and maintenance of synapses (Waites et al. 2005; Ziv and Garner 2004). In the case of the axonal transport of SV proteins, anterograde movement is driven by the kinesin motors, KIF1A (Okada et al. 1995) and KIF5B (Nakata and Hirokawa 2003; Song et al. 2009), and retrograde transport is mediated by dynein (Fejtova et al. 2009; Paschal and Vallee 1987; Schnapp and Reese 1989). However, how SV proteins are sorted to presynaptic boutons remains unclear.

A great deal of progress has been made concerning the postal system by which proteins are selectively sorted to dendrites. A direct pathway, with proteins traveling directly from the soma to dendrites, has been shown to be established early in neuronal development (Burack et al. 2000; Karasmanis et al. 2018; Petersen et al. 2014; Silverman et al. 2001). Studying the sorting of axonal cargo, specifically SV proteins, in mature neurons has proven to be more challenging, in part due to the limited flux of materials to neurites after synaptogenesis, a point we return to below. In mammalian neurons, two SV protein trafficking pathways have been proposed: the selective retention model posits that there is non-polarized delivery of transport vesicles to axons and dendrites, followed by the retention of protein in axons and preferential endocytosis from the dendritic compartment and subsequent re-routing to axons (Fletcher-Jones et al. 2019; Sampo et al. 2003). The second pathway is a variant of this model. It differs in that axon-destined transport vesicles move through axons and dendrites but do not fuse with the somatodendritic plasma membrane (PM) (Burack et al. 2000; Nabb and Bentley 2022). This second model has been termed ‘direct transport’, despite the observation that transport vesicles carrying axonal proteins often entered dendrites, to emphasize the lack of fusion of these vesicles in the somatodendritic domain. After decades of research, the widespread conclusion is that SV proteins are trafficked with a low degree of selectivity into both axons and dendrites of mammalian neurons, and are either selectively retained in, or have partially biased transport towards, axons (Bentley and Banker 2016). In the current study we address a third model which theorizes that SV proteins are selectively delivered to axons, without entering dendrites, as indicated in *C. elegans* DA9 bipolar neurons and pseudounipolar dorsal root ganglion cells (Gumy et al. 2017; P. Li et al. 2016) Importantly, direct and selective transport of SV proteins to axons has not been clearly demonstrated in mammalian neurons with classically polarized axonal and dendritic arbors (Karasmanis et al. 2018; Nakata and Hirokawa 2003).

We emphasize that because SVs are a hallmark of presynaptic boutons and represent a striking specialization within axons, the trafficking of SV proteins to nerve terminals is often used as a proxy for axonal transport. Moreover, how these proteins are incorporated into SVs once they make their way into axons remains completely unknown. Again, these gaps in knowledge are due, in part, to the low rate of SV protein egress from the soma once neurons mature and switch from establishing to maintaining their polarity. This makes it difficult to monitor movement of SV precursors via live-cell imaging, without overexpressing the protein of interest.

The main objective of the current study is to trace the path that two distinct SV proteins take from the soma to axons in mammalian neurons. We focused on synaptotagmin (SYT) 1, a Ca^2+^ sensor that regulates rapid neurotransmitter release (Chapman 2008), and SYB2, a vesicular (v-) SNARE protein, also known as VAMP2, that assembles into *trans*-SNARE complexes to catalyze membrane fusion (Südhof and Rothman 2009). We note that SYB2 has been used in axonal transport studies for decades, and was included here to directly address the idea that its polarized distribution is achieved via either of the two non-polarized delivery models outlined above (Nabb and Bentley 2022; Sampo et al. 2003). Additionally, these two SV proteins were selected because SYT1, a canonical type I transmembrane protein, is co-translationally inserted into the endoplasmic reticulum (ER) (Perin et al. 1990; Shao and Hegde 2011), whereas SYB2, a type II tail-anchored protein, is post-translationally inserted into the ER (Kutay et al. 1995). Moreover, SYT1 and SYB2 have been shown to be trafficked on different kinesin motors, KIF1A (Okada et al. 1995) and KIF5B (Nakata and Hirokawa 2003; Song et al. 2009), respectively. The selection of two distinct SV proteins with different topologies, and consequently different biosynthetic pathways, as well as different trafficking motors, enabled us to investigate whether there is a conserved mechanism underlying polarized transport. More generally, using these SV proteins as proxies for axonal cargo, we address the widely accepted model that, in mammalian neurons, axonal proteins make their way into all neurites, but are eventually selectively retained in axons.

To overcome the technical challenges discussed above, we took advantage of recently developed tools to sequester SV proteins in the ER of mature neurons. We then released them in a synchronized manner, after synaptogenesis, to track their path to synapses after leaving the Golgi (Boncompain et al. 2012). We combined this system with a HaloTag labeling approach (Grimm et al. 2015; Grimm and Lavis 2021; Los et al. 2008) to follow the fate of SYT1 and SYB2 as they leave the soma and are ultimately delivered to nerve terminals. In these experiments, careful attention was paid to expression levels since overexpression results in the spillover of SV proteins into inappropriate compartments, thus obscuring polarized transport (Pennuto 2003). These experiments uncovered a novel pathway in which transport vesicles, bearing SV proteins, are directly and selectively delivered to axons. Furthermore, upon delivery, these vesicles fuse with the presynaptic PM creating a hub, or reservoir, from which SVs are generated. These findings argue against the selective retention model and other non-polarized transport pathways; rather, SV proteins undergo direct egress from the soma to axons without entering dendrites.

## Results

### Using RUSH to study egress of SV proteins from the soma of cultured neurons

We took advantage of the retention using selective hooks (RUSH) system to study the sorting itinerary of newly synthesized SV proteins (**Fig. 1A**) (Boncompain et al. 2012). In the RUSH system, proteins of interest are retained in the ER and released upon addition of a small molecule. The ability to synchronize protein release from the ER makes it possible to observe their trek to their target destination. This system was chosen over other rapalog-induced dimerization systems, such as FKBP12, because acute application of the small molecule induces release. In contrast, other approaches require the chronic application of a small molecule to maintain the retention of protein. We also avoided optical-based retention/release approaches due to the potential for premature cargo release, due to light exposure during culture maintenance and imaging. Retention of the SV proteins, SYT1 and SYB2, was accomplished by appending a streptavidin-binding peptide (SBP) to the intravesicular end of each protein; translocation of the amino-terminus of SYT1 into the ER was aided by the addition of a pre-prolactin leader sequence (**Fig. 1A, 1B**). The tagged proteins bind to the co-expressed streptavidin ‘hook’, which is localized to the ER by a retention signal (Lys·Asp·Glu·Leu; KDEL), thus retaining them. Addition of the small molecule, biotin, displaces the SBP and allows natural egress to occur. Each cargo also includes a HaloTag that was used for visualization.

**Fig. 1.**
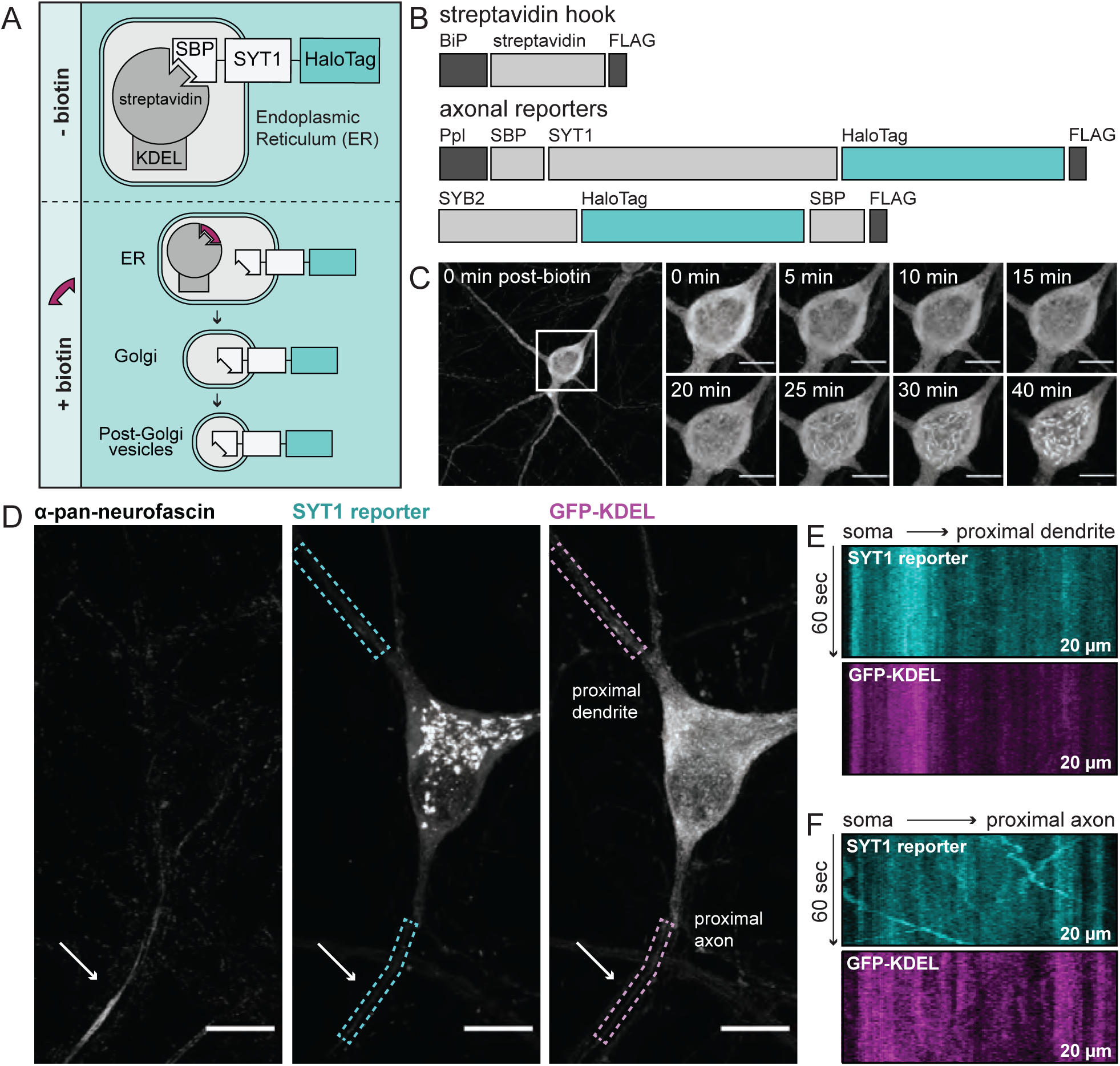
Using RUSH to study egress of SV proteins from the soma of cultured rat hippocampal neurons. (**A**) A cartoon of RUSH; pre- and post-biotin conditions are shown. (**B**) Schematic of the streptavidin hook, and SYT1 and SYB2 reporter RUSH constructs: BiP, a signal peptide that drives translocation into the ER; FLAG, provides a means to detect each construct; SBP, streptavidin binding peptide; Ppl, a pre-prolactin leader sequence to translocate the SBP into the ER. In all cases the reporter is a HaloTag. (**C**) Representative super-resolution fluorescent live-cell MAX projection images from rat neurons at 15 days *in vitro* (DIV). Images of SYT1 reporter immediately after biotin addition with enlarged insets to detail the time-course of release (since SYT1 and SYB2 behaved similarly, only SYT1 images are shown in panels (**C-F**). Inset scale bar is 10 μm in all panels. (**D**) Image of a neuron expressing the streptavidin hook, SYT1 reporter, and ER-targeted GFP, 30 minutes after biotin addition. Live-cell labeling with an anti-pan-neurofascin antibody was used to identify the axon initial segment (AIS; arrow); dendrites were identified by exclusion and morphology. SYT1 was labeled with JF549 HaloTag ligand, and kymographs of this reporter, along with GFP-KDEL, were generated from the regions indicated by dashed boxes (20 μm long). Kymographs from a proximal dendrite (**E**) and proximal axon (**F**) are shown.

It is known that overexpression can cause SV proteins to mislocalize to other compartments, especially the PM (Pennuto 2003). To mitigate this confound, the viruses used to express SYT1 and SYB2 were carefully titrated to achieve a sparse transduction such that only a select few neurons were expressing minimal levels of the tagged protein. To further ensure low levels of expression, cells that had lower than average fluorescence were selected for imaging. Within this low expression paradigm, live-cell imaging and immunocytochemistry (ICC) confirmed the localization of the SV reporter proteins within the ER prior to biotin-triggered release, and then with the Golgi and eventual endpoint targeting to synapses, following release (**Fig. 1C, Supplementary Fig. S1**).

With the temporal control afforded by this assay, we can designate an exact starting position and time of release in the cell, and record trafficking events immediately upon exit from the Golgi. To definitively identify axons, we used an extracellular pan-neurofascin antibody to label the initial segment of live neurons (**Fig. 1D**). Upon release of ER-tethered SYT1, we observed little to no transport activity in proximal dendrites as shown by the lack of diagonal lines in the kymograph (**Fig. 1E**); however, there was noticeable movement of tagged SYT1 as it began to egress from the soma directly into axons in an anterograde direction, as represented in the kymograph by diagonal lines with a negative slope (**Fig. 1F**). These initial findings contradict the idea of SV proteins being trafficked with low selectivity into axons and dendrites (Nabb and Bentley 2022; Sampo et al. 2003), and warrant a deeper examination. We highlight that the majority of anterograde-moving vesicles observed in proximal axons did not contain the co-expressed ER-targeted GFP (**Fig. 1F**), confirming that these are post-ER organelles. In contrast, the stationary SYT1 signal in the proximal dendrites colocalized with ER-targeted GFP (**Fig. 1E**), indicating the observed signal is protein in the ER, rather than post-Golgi transport vesicles. Taken together, the lack of movement in proximal dendrites, and the robust anterograde trafficking in axons, suggest the existence of a selective and specific pathway that targets SYT1 to presynaptic boutons.

### A direct and selective axonal transport pathway for SYT1 and SYB2

To explore this seemingly novel trafficking pathway, we expanded our experiments to include a second SV protein, SYB2, and to analyze transport in both proximal and distal compartments of both axons and dendrites. Proximal compartments were defined as the first 20 μm of a neurite as it emerges from the cell body. Distal compartments were defined as a secondary branching of a dendrite or, for axons, a distance of ∼150 microns from the soma, which is beyond the axon initial segment (**Fig. 2A**). With these selection criteria, transport was quantitatively assessed by generating and analyzing kymographs of post-Golgi vesicles carrying the SV protein of interest (**Supplementary Fig. S2**). As was first seen in **Fig. 1**, we again observed robust transport of the SYT1 reporter in proximal axons and extended these observations to distal axons; proximal and distal dendrites had little to no detectable trafficking of SYT1-containing transport vesicles (**Fig. 2B-D, Supplementary Fig. S3A**). This trend was also observed for the SYB2 reporter, allowing us to generalize our findings to topologically distinct SV proteins that are carried by different kinesin motors (**Fig. 2E-G, Supplementary Fig. S3B**). Furthermore, all neurites were observed for the exact same duration, so the volume of transport activity in each compartment can be directly compared, further supporting the idea that there is minimal transport of SV proteins in dendrites. Of the transport vesicles observed, both SYT1 and SYB2 reporters were preferentially transported in the anterograde direction in axons (**Fig. 2H, 2I**). Again, few, if any, puncta were observed in dendrites (**Fig. 2J, 2K**). These data confirm a general trafficking pathway that—in contrast to previous models (Nabb and Bentley 2022; Sampo et al. 2003)— does not include significant flux through dendrites. Rather, our findings establish a transport pathway that selectively and specifically routes SV proteins to axons. We cannot rule out that low levels of transport through dendrites likely occur below the limit of detection in our system, but the predominant mode of transport was observed to be direct and specific to axons.

**Fig 2.**
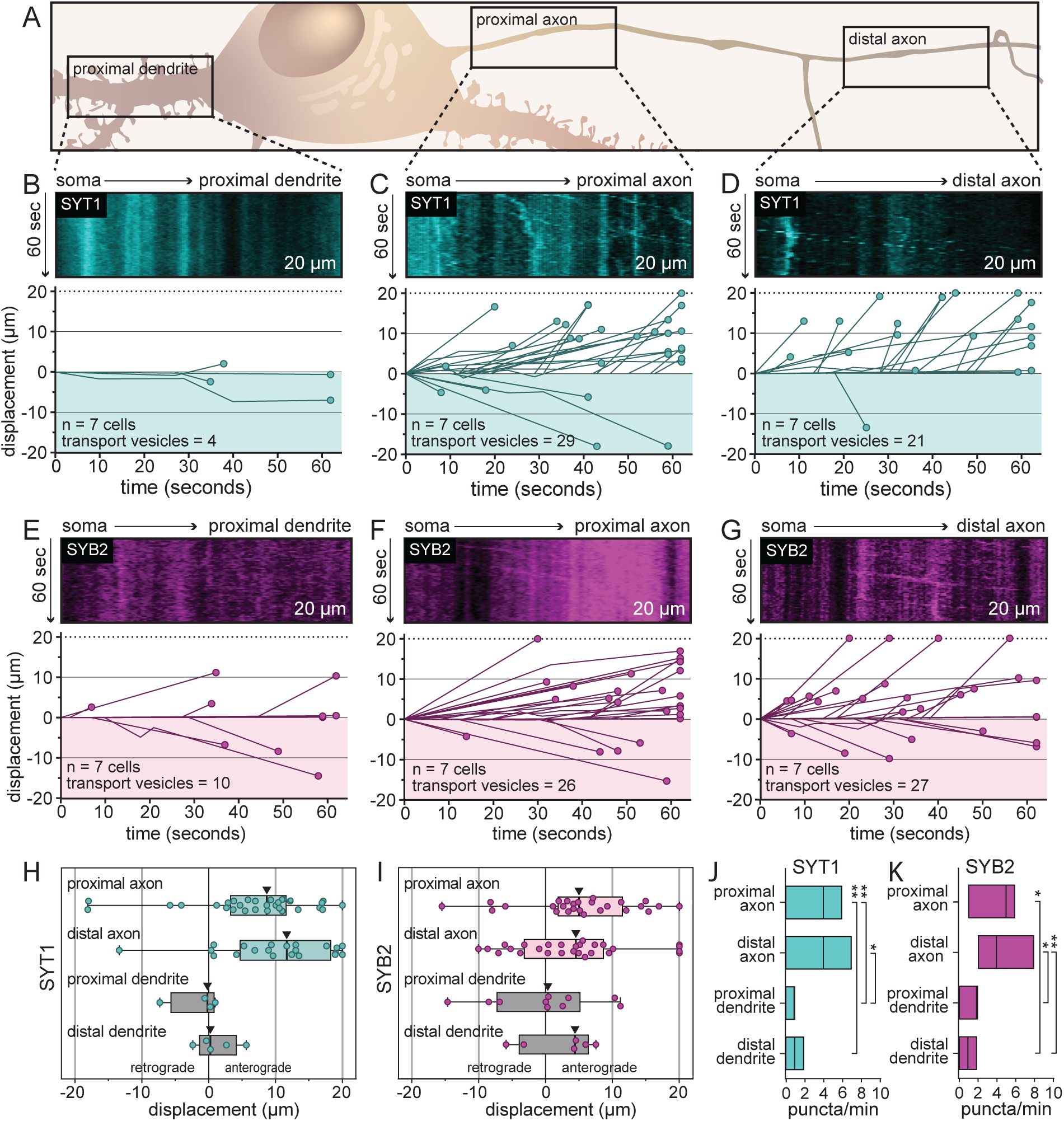
A direct and selective axonal transport pathway for SYT1 and SYB2 in rat hippocampal neurons. **(A)** llustration outlining the proximal and distal regions that were imaged for each neuron. (**B**) Representative kymographs from the proximal dendrite of rat hippocampal neurons at 14-16 DIV after release of the tethered SYT1 reporter, revealing an absence of SYT1-bearing mobile organelles. For panels (**B-G**), all data were quantified and plotted immediately below the kymographs; the number of cells and tracks are also indicated.(**C**,**D**) Representative kymographs from proximal and distal axons showing robust movement of the released SYT1 reporter, suggesting a direct axonal trafficking pathway. (**E-G**) Same as for panels (**B-D**) but using neurons expressing the SYB2 reporter. Displacement of transport vesicles containing the SYT1 (**H**) or SYB2 reporters (**I**) is plotted in the anterograde (positive) or retrograde (negative) direction with respect to the soma; arrowheads indicate median values. Both SV proteins are primarily trafficked in the anterograde direction. mean values and descriptive statistics are found in **Supplemental Table S1**. (**J**) Flux of the SYT1-bearing transport vesicles, in the indicated compartments, are presented as mean values <u>±</u> SD. Data were collected from 7 cells. A one-way ANOVA with multiple comparisons was run; p-values were as follows: proximal axon vs. distal axon = 0.563; proximal axon vs. proximal dendrite = 0.00210; proximal axon vs. distal dendrite = 0.00470; distal axon vs. proximal dendrite = 0.0463; distal axon vs. distal dendrite = 0.0914; proximal dendrite vs. distal dendrite = 0.987. (**K**) Same as (**J**), but for the SYB2 reporter. Data were collected from 7 cells. Statistical tests were run as in (**J**); and p-values were as follows: proximal axon vs. distal axon = 0.998; proximal axon vs. proximal dendrite = 0.0584; proximal axon vs. distal dendrite = 0.0130; distal axon vs. disimal dendrite = 0.0408; distal axon vs. distal dendrite = 0.00880; proximal dendrite vs. distal dendrite = 0.907. Mean values and descriptive statistics are found in **Supplemental Table S1**.

To increase rigor, we conducted control experiments to determine whether our assay can, in fact, accurately identify dendritic transport. For this, we chose the transferrin receptor (TfR), a protein localized to dendrites (Burack et al. 2000; H. Li et al. 2016; West et al. 1997a). We observed that TfR was overwhelmingly trafficked to dendrites without passing through axons, even when purposefully overexpressed (**Supplementary Fig. S4**). This demonstrates that our assay can, indeed, reveal selective and direct dendritic transport, further validating our findings of a direct transport pathway for SYT1 and SYB2.

### The direct and selective transport of SV proteins is obscured by overexpression

As stated in the Introduction, the expression levels of SV proteins can affect their localization; namely, overexpression results in spillover of these proteins into other compartments, including the PM (Marks et al. 1996; Pennuto 2003). Additionally, catch-and-release assays can also cause mislocalization by overwhelming the trafficking machinery of the early secretory pathway upon the bulk release of protein (Adams et al. 2019). Indeed, in agreement with previous studies, when we drastically overexpressed SYT1 via transfection, it spilled over into dendrites and ‘coated’ the PM of all neurites; it also caused the growth of filopodia-like structures in the somatodendritic domain (**Fig. 3A**) (Feany and Buckley 1993).

**Fig. 3.**
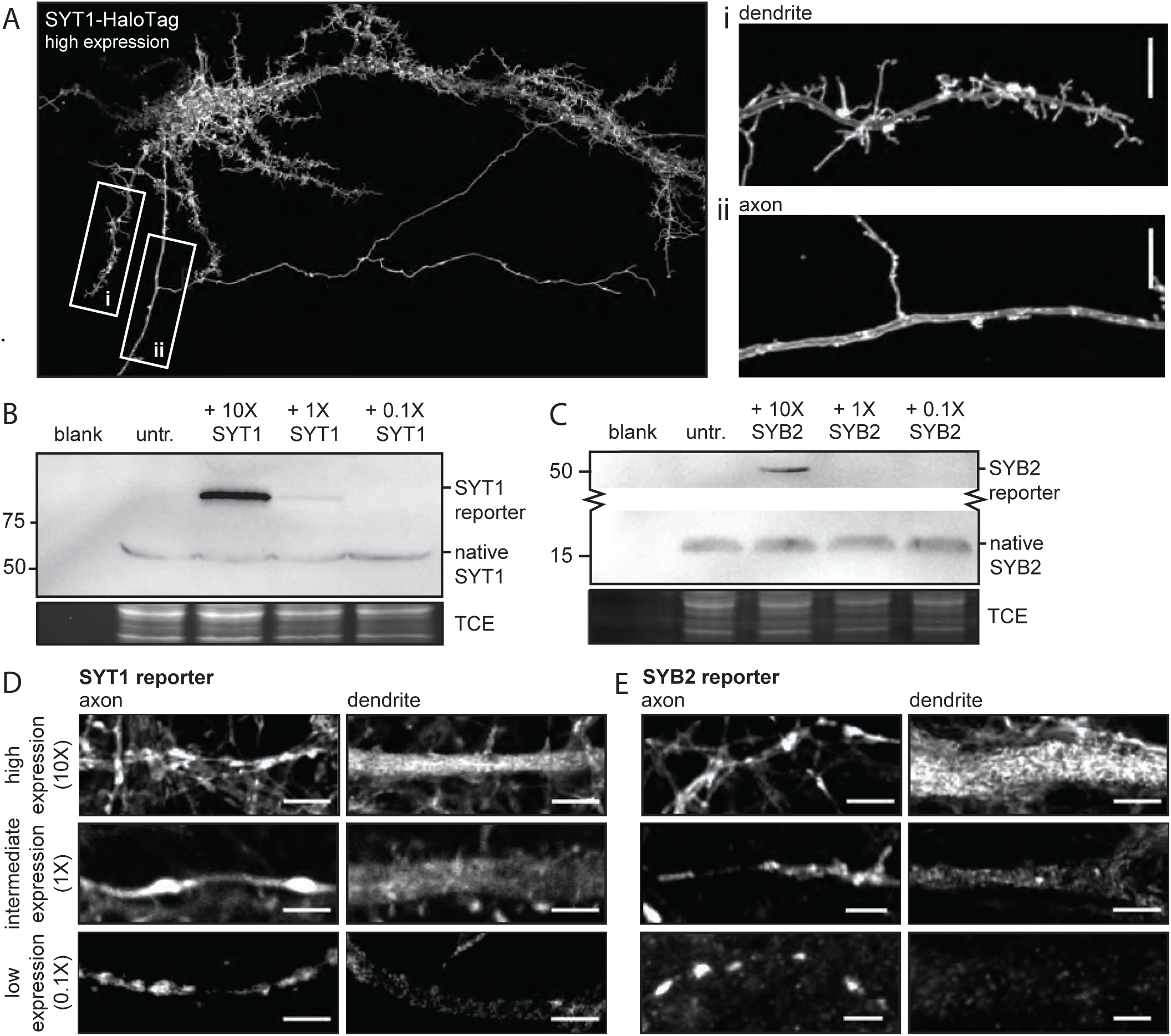
The direct and selective transport of SV proteins is obscured by overexpression. (**A**) A super-resolution MAX-projection image of 15 DIV rat neurons expressing SYT1-HaloTag at high levels, as evidenced by the localization of SYT1-HaloTag in both axons and dendrites, along with the formation of filopodia-like structures in the somatodendritic compartment. The construct is visualized with JF549 HaloTag ligand. The boxed regions were expanded to show the overexpressed protein is present in both the dendritic (i) and axonal (ii) PM. Scale bar is 5 μm. (**B**) An immunoblot of 15 DIV rat neurons expressing the SYT1 reporter at low, intermediate, and high levels using virus. Probing with an αSYT1 antibody reveals the exogenously expressed SYT1 reporter is present at much higher levels than the endogenous protein, confirming high expression. (**C**) Likewise, an immunoblot of rat neurons expressing the SYB2 reporter at various levels was probed with an αSYB2 antibody to reveal the degree of overexpression compared to the native protein. Lower expression levels of SYT1 and SYB2 escaped detection due to the sparse transduction. Super-resolution optical sections of the SYT1 (**D**) and SYB2 (**E**) reporters at low, intermediate, and high expression levels, in axons and dendrites, demonstrate that as expression levels increase, SV reporters spillover into dendrites. Scale bar represents 2.5 μm. Corresponding axon and dendrite images at each expression level were adjusted with the same linear brightness and contrast settings.

To formally address these concerns, the SYT1 and SYB2 RUSH reporter constructs were expressed at high, intermediate, and low levels, by controlling the amount of virus used for transduction. Western blot analysis confirmed the degree of overexpression of each protein, as compared their endogenous counterparts (**Fig. 3B, 3C**). We note that, at lower virus titers, drastically fewer cells were transduced, so the exogenously expressed proteins escaped detection. This lowest virus titer was used for experiments conducted in **Fig. 1** and **Fig. 2**. At high expression levels, both the SYT1 and SYB2 reporters appeared throughout cells at steady state (**Fig. 3D, 3E**). A similar trend was observed at intermediate expression levels, albeit with lower signals in dendrites. Only when expression levels were low—indeed, further dilution of the virus resulted in a lack of transduced cells —and the dimmest cells were selected, did we observe the polarized, and pre-synaptic distribution of the SYT1 and SYB2 reporters that is characteristic of SV proteins (**Fig. 3D, 3E**) (Chapman 2008; Südhof and Rothman 2009). We note that, under this low expression paradigm, neurons are quite dim and, thus, challenging to image (i.e., required sensitive microscopy and a concentrated release of protein via the RUSH assay to observe transport), which may explain why higher expression levels are often employed in axonal transport studies.

Exogenously expressed SV proteins spillover into dendrites at high, and even intermediate, expression levels at steady state (**Fig. 3A, 3D, 3E**), likely due to indiscriminate transport when expression levels are not carefully controlled. Therefore, it is imperative to use a low expression paradigm to study the native transport of these proteins. We note that a similar trend of protein mislocalization at high expression levels was observed using a transfection approach (data not shown); only when we “diluted” the plasmid of interest, by mixing and co-transfecting it with a dummy plasmid, were we able to minimize overexpression artifacts. Simply reducing the total amount of the plasmid of interest for transfection was not sufficient to mitigate the rampant mistargeting. Taken together, these data demonstrate the extent to which overexpression can cause SV proteins to be mistargeted and ultimately obscure their native transport pathway. Additionally, these results help to reconcile the discrepancies between the current study and previous studies reporting the trafficking of proteins like SYB2 in both axons and dendrites (Nabb and Bentley 2022; Sampo et al. 2003).

### Molecular determinants that underlie the polarized transport of SYT1 to axons

The sorting motifs in SYT1 are not well defined. However, previous work suggested a role for post-translational modifications in its localization, namely palmitoylation (Chapman et al. 1996; Heindel et al. 2003; Kang et al. 2004) and glycosylation (Atiya-Nasagi et al. 2005; Han et al. 2004; Kwon et al. 2012; Perin et al. 1991). To address this, all five putative palmitoylation sites and all three glycosylation sites of SYT1 were mutated to prevent these modifications (**Fig. 4A**). Hereafter, this mutant is referred to as the SYT1 palmitoylation/glycosylation mutant, or SYT1-PGM. In parallel, we assessed the roles of the tandem C2-domains of SYT1, which sense Ca^2+^ and interact with a variety of effectors, for targeting to nerve terminals. We note that some degree of synaptic localization of SYT1 was preserved when individual C2 domains were deleted; however, deletion of both domains caused the truncated protein to be marooned on the PM (Courtney et al. 2019). How this deletion mutant was sorted and transported remains unknown, so this mutant, termed SYT1ΔC2AB, was also analyzed using the RUSH assay (**Fig. 4A**). All experiments were done in a SYT1 knockout background to avoid potential homomeric interactions with endogenous SYT1 (Brose et al. 1992; Courtney et al. 2021; Perin et al. 1991), so that mutant protein cannot ‘piggyback’ onto the native protein in the secretory pathway and obscure potential transport defects associated with the mutant protein (**Fig. 4B**).

**Fig. 4.**
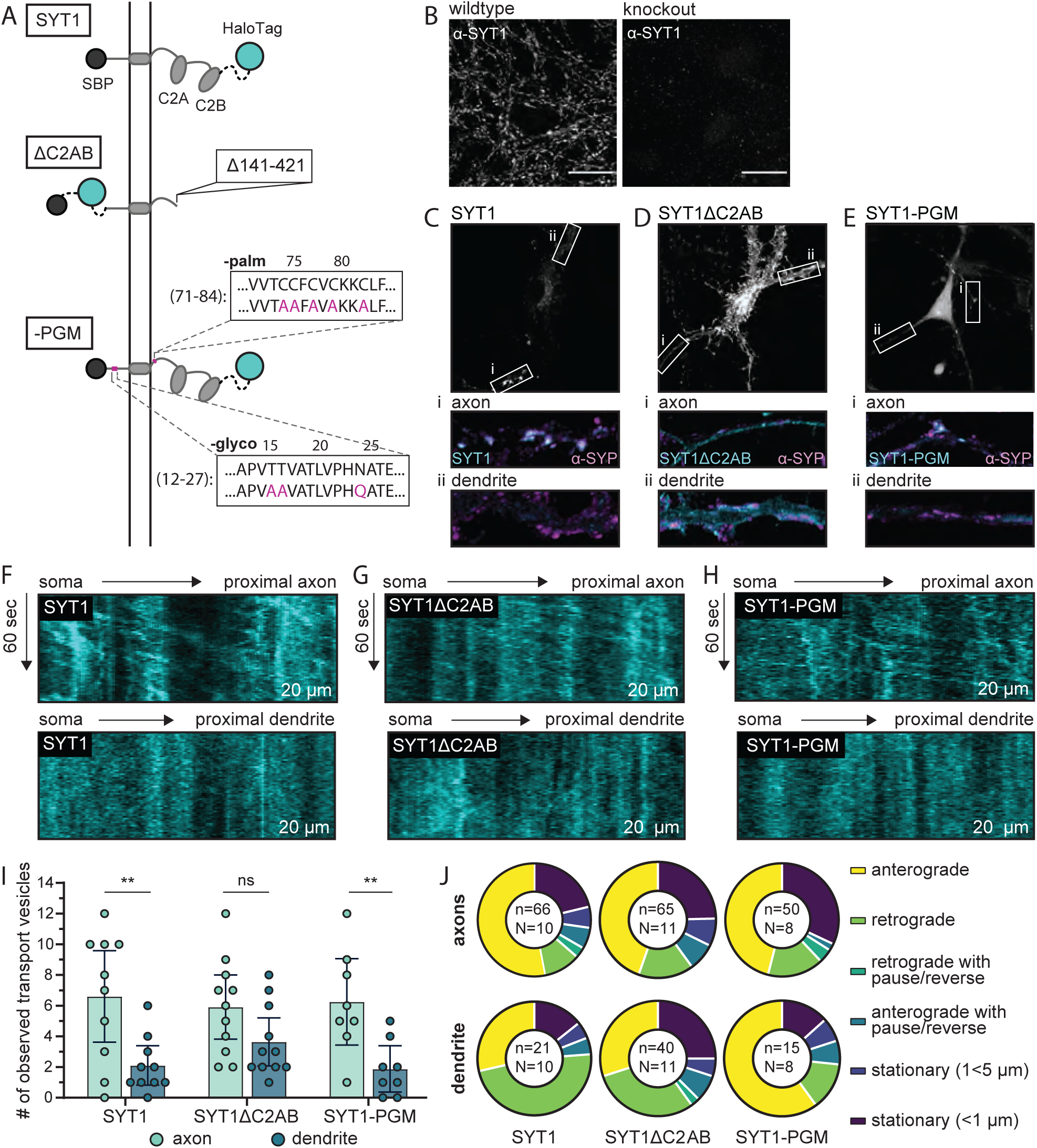
Molecular determinants that underlie the polarized transport of SYT1 to axons in mouse hippocampal neurons. (**A**) Illustration of RUSH reporters used for these experiments: WT SYT1 reporter, SYT1 truncated after position 140 (SYT1ΔC2AB), and the SYT1 palmitoylation and glycosylation mutant (SYT1-PGM). Each construct has a HaloTag for visualization. (**B**) ICC confirms the knockout of endogenous SYT1. For WT and knockout conditions, identical laser and gain settings were used. Linear brightness and contrast adjustments were applied to both conditions. (**C-E**) The endpoint localization of WT SYT1, SYT1ΔC2AB, and SYT1-PGM was visualized by labeling the appended HaloTag with JF549. The boxed regions were expanded and are shown below each panel to better reveal the localization of each construct in axons (i) and dendrites (ii), as compared to the α-SYP ICC signals. Note that all neurons were immunostained for SYP, but only a handful of cells expressed each SYT1 construct. ICC images were adjusted to the brightest area of the image to aid in visualization. Brightness and contrast settings were kept consistent between corresponding axon/dendrite insets. In all cases, images were adjusted with linear brightness and contrast. Representative kymographs from proximal axons showing robust anterograde movement of the released SYT1 (**F**), SYT1ΔC2AB (**G**), and SYT1-PGM (**H**) reporters as compared to dendrites, demonstrating selective trafficking in axons. (**I**) The number of transport vesicles was plotted for each construct as the mean with 95% CI, and differences between axons and dendrites were analyzed for each construct using an unpaired t-test (p_SYT1_ = 0.00570; p_ΔC2AB_ = 0.0670; p_-PGM_ = 0.00600). Data were collected for 10 cells (SYT1), 11 cells (SYT1ΔC2AB), or 8 cells (SYT1-PGM) from 4 litters. Mean values and descriptive statistics are found in **Supplementary Table S2**. (**J**) The movement of each transport vesicle categorized as anterograde, retrograde, retrograde with pause/reverse, anterograde with pause/reverse, stationary (1<5 μm), or stationary (<1 μm) and plotted as a fraction of the total number of transport vesicles observed for each compartment, for each construct. The total number of (n) transport vesicles from (N) cells are indicated. Exact fractions can be found in **Supplementary Table S2**.

To begin, the endpoint targeting of each reporter construct was established at low expression levels. The full-length SYT1 reporter was included as a positive control, and was correctly targeted to synapses, as confirmed by its colocalization with the synaptic marker, synaptophysin (SYP) (**Fig. 4C**). The truncated SYT1 protein, SYT1ΔC2AB, was present throughout axons, likely in the PM (Courtney et al. 2019), and was also observed in the somatodendritic compartment, indicating mistargeting of the protein (**Fig. 4D**). The SYT1-PGM construct accumulated in the axonal compartment, where it was co-localized with SYP, and was not detected in dendrites, similar to our findings using the wildtype (WT) SYT1 reporter (**Fig. 4E**).

We then studied the transport of the PGM and ΔC2AB mutants in the RUSH assay. Each construct was successfully sequestered in the ER and released. Notably, all constructs successfully left the Golgi in transport vesicles and did not immediately fuse with the somatic PM, but instead were trafficked into neurites. We note that these experiments were conducted without the addition of ER-targeted GFP, because the RUSH assay workflow improved when cells only expressed the hook and the reporter (i.e., constructs released more reliably, and the post-Golgi vesicles were brighter and easier to visualize).

In knockout neurons, the WT SYT1 reporter was—again—trafficked in a polarized manner to axons (**Fig. 4F, 4I**). In contrast to previous studies (Han et al. 2004; Heindel et al.2003; Kang et al. 2004; Kwon et al. 2012), but consistent with our findings under steady state, the transport of SYT1-PGM was virtually the same as the WT protein: it was specifically trafficked to axons (**Fig. 4H, 4I**). SYT1ΔC2AB behaved differently; this construct entered axons and dendrites at similar rates, with a non-significant trend toward preferential entrance into axons (p=0.0670) (**Fig. 4G, 4I**).

The movements of each transport vesicle in axons and dendrites, for each of the three constructs, were quantified and plotted (**Fig. 4J**, see also **Table S2**). All three constructs tended to proceed in an anterograde direction in axons. In dendrites, the majority of transport vesicles carrying SYT1 and SYT1ΔC2AB moved in a retrograde direction, though a considerable fraction of SYT1ΔC2AB vesicles were stationary. Surprisingly, SYT1-PGM overwhelmingly moved in an anterograde direction in dendrites under the non-equilibrium conditions of these experiments. However, the total number of transport vesicles carrying SYT1 and SYT1-PGM in dendrites was relatively low, so the observed differences should be interpreted with caution. Taken together, these experiments reveal that the tandem C2 domains play a role in the proper targeting of SYT1. In contrast, palmitoylation and glycosylation were dispensable for selective targeting of SYT1.

### Transport vesicles deliver SYT1 to the presynaptic plasma membrane, creating a depot for SV biogenesis

Finally, we sought to visualize the endpoint destination of newly delivered SYT1 within synapses. Previous studies have established that SVs are assembled at the pre-synapse (Buckley et al. 2000a; Nakata et al. 1998; Okada et al. 1995; West et al. 1997b), but how they are first generated remains unknown. It has long been hypothesized that SV proteins, prior to their incorporation onto nascent SVs, are first trafficked to the presynaptic PM as the final destination of their maiden voyage to synapses (Buckley et al. 2000b; Feany et al. 1993; Hannah et al. 1999; Regnier-Vigouroux et al.1991); however, this idea stems from experiments done in PC12 cells and CHO fibroblasts, which do not contain SVs. Since SYT1 is present in the PM at steady state (Sankaranarayanan et al. 2000), and overexpressed SYT1 also accumulates in the PM (**Fig. 3**), it is reasonable to postulate that this SV precursor is initially trafficked through the PM of mammalian neurons as a necessary part of its life cycle.

We addressed this longstanding question by developing a novel HaloTag labeling approach to conduct pulse chase experiments using permeant and non-permeant Janelia Fluor (JF) HaloTag ligands (HTL) (Grimm et al. 2015). To assess whether SYT1 is first delivered to the presynaptic PM in mammalian neurons, we appended a HaloTag to its luminal domain (HaloTag-SYT1) so that the tag is exposed to the outside of the cell when SYT1 is in the PM (**Fig. 5A**); this construct was sparsely co-transfected with a synaptophysin-GFP (SYP-GFP) fusion protein, to mark synapses. The ratio of SYT1 to SYP plasmid in the co-transfection was experimentally determined so that the HaloTag-SYT1 could be visualized, but was minimally expressed via concurrent dilution with the SYP plasmid. Immediately after co-transfection, neurons were grown with or without JF549i, a non-permeant fluorescent HTL, in the culture media so that any tagged SYT1 that passed through the PM would be labeled there and, as a result, visualized (Xie et al. 2017). Impermeability of the ligand, under our experimental conditions, was confirmed empirically (**Supplementary Fig. S5**). After 6 days the degree of labeling with JF549i was assessed via imaging, and the neurons were subsequently challenged using a permeant ligand, JF549, that has nearly the same structure, and identical fluorescence properties, as JF549i. This subsequent challenge labels, and hence exposes, any remaining unlabeled protein that did not pass through the PM. This labeling scheme is illustrated in **Fig. 5B, 5C**.

**Fig. 5.**
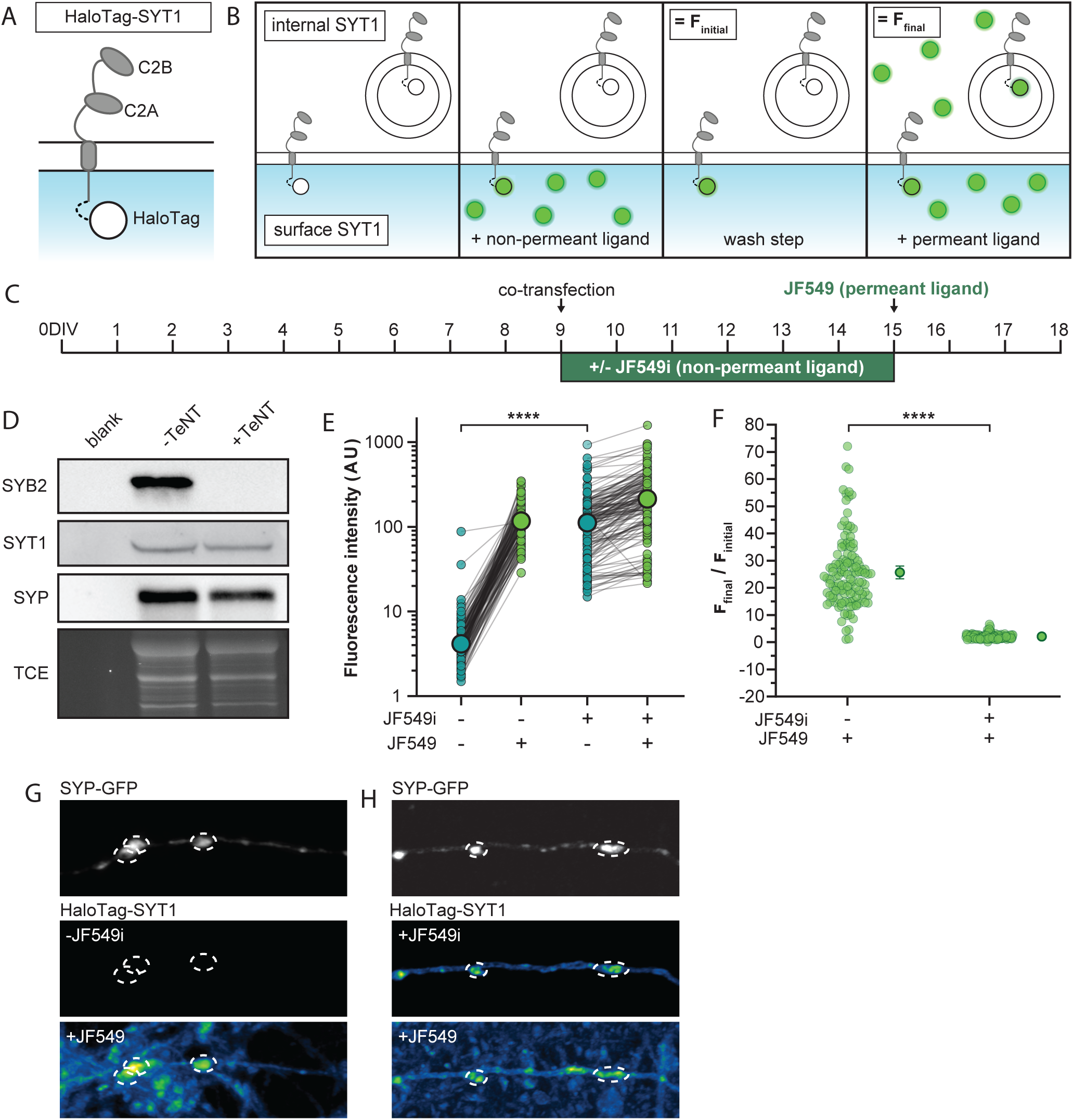
Transport vesicles deliver SYT1 to the presynaptic plasma membrane, creating a depot for SV biogenesis. (**A**) Illustration of SYT1 with a luminal HaloTag (HaloTag-SYT1) to allow for selective labeling at the PM. (**B**) Schematic of the HTL labeling protocol starting with the non-permeant ligand (JF549i) incubation step to label surface protein, washing out the ligand and labeling with the permeant ligand (JF549) to label all protein. F_initial_ is the signal from labeling with the non-permeant ligand, and F_final_ is the subsequent signal after labeling with permeant ligand. (**C**) Timeline for the transfection and labeling protocols. Cultured rat hippocampal neurons were co-transfected on 9 DIV with HaloTag-SYT1 and SYP-GFP; the latter construct was included to mark synapses and ‘dilute’ the HaloTag-SYT1 plasmid to achieve lower expression levels. Half of the coverslips were incubated in non-permeant HTL immediately after co-transfection to label any tagged protein that was delivered to the PM. Six days later (15 DIV) neurons were rinsed, imaged, and incubated with permeant ligand, to label any remaining tagged protein, and imaged again. (**D**) Immunoblot of cells transduced with a virus to express the light chain of TeNT, resulting in the cleavage of SYB2 and the inhibition of SV recycling. Blots were probed for SYB2, SYT1 and SYP, with a TCE loading control. (**E**) The fluorescence intensity of HaloTag-SYT1 labeling at the PM, for cells grown with and without JF549i (before and after permeant ligand addition) for 6 days, was quantified and plotted. Enlarged data points indicate the median values. The degree of labeling at the plasma membrane by the JF549i HTL, prior to permeant ligand addition, was analyzed with a Mann-Whitney test; p-value = <0.0001. (**F**) The data from (**E**) were replotted as the normalized (F_final_/F_initial_) change in fluorescence upon adding a permeant fluorescent ligand to cells grown with or without non-permeant ligand for 6 days; mean values with 95% CI are plotted. Data were analyzed with a Mann-Whitney test; p-value = <0.0001. Panels (**E, F**) show data from 156 synapses cultured in the presence of JF549i, and 136 synapses grown in the absence of this HTL. Data for both groups were from 8 fields of view from 4 different litters. Mean values and descriptive statistics can be found in **Supplementary Table S3**. Panels (**G, H)** are representative, super-resolution images of SYP-GFP to mark synapses (dashed lines), and the HaloTag-SYT1 signals under the indicated conditions; in the bottom panels the JF549 ligand was not washed away, resulting in a higher background. For all conditions, identical laser and gain settings were used. Any linear brightness and contrast adjustments were applied to all conditions.

Synaptic activity in our cultures, and hence the recycling of SVs, could contribute to SYT1 passing through the PM. To minimize this potential confound, the light chain of tetanus toxin (TeNT) was co-expressed, using lentivirus, to cleave endogenous SYB2 and inhibit synaptic activity and SV recycling (**Fig. 5D**) (Bao and Das et al. 2018; Schiavo et al. 1992). This ensures that any labeling of SYT1 that occurred at the PM was largely independent of the SV cycle. Cleavage of SYB2 by TeNT did not affect the expression of other canonical SV proteins (**Fig. 5D**).

To quantify labeling of HaloTag-SYT1 at the synaptic PM, co-transfected SYP-GFP was used to define individual synapses; fluorescence intensity was measured at each synapse before and after the addition of the permeant HTL, JF549. As expected, in cultures grown in the absence of JF549i, we observed a dramatic increase in the fluorescence intensity at synapses upon the addition of JF549. However, synapses cultured with JF549i for 6 days exhibited minimal changes in fluorescence after addition of the permeant JF549. These findings reveal that the majority of the tagged SYT1 was already labeled at the PM (**Fig. 5E**). These data were replotted as the change in fluorescence upon addition of the permeant JF549 dye, further illustrating the negligible change in fluorescence at each synapse for cultures grown in the presence of JF549i (**Fig. 5F**). This analysis quantitatively shows that the majority of SYT1 passes through the PM, independent of significant levels of synaptic activity. We emphasize that these results were readily apparent by eye, as shown in the representative images of cultures grown with and without the non-permeant ligand and chased with a permeant ligand (**Fig. 5G, 5H**). We conclude that newly delivered SYT1 is efficiently incorporated in the PM, creating a depot from which SV biogenesis can occur, potentially via on-going synaptic activity.

## Discussion

A neuron’s ability to sort proteins and transport cargo to synapses underlies the function of the nervous system and is a process that is maintained throughout the lifespan of the cell. As such, several theories have been proposed for how transport occurs. The leading model, for the past two decades, has been that axonal proteins are delivered indiscriminately to all neurites and are subsequently selectively retained in axons (Bentley and Banker 2016; Sampo et al. 2003). In sharp contrast, the transport of dendritic cargos has been shown to be selective, and vesicles carrying dendritic cargo are trafficked directly to dendrites without entering axons. This presents an apparent paradox, because both dendritic and axonal arbors can have elaborate morphologies, and transporting the cargos destined for axons through another exceptionally complex compartment complicates this sorting process. Consequently, selective retention, and other variants of this non-selective transport theory, appeared to be a high-effort, low-reward method of establishing and maintaining polarity. While inefficient mechanisms cannot be ruled out, the development of a specific postal system for dendrites, but not axons, remained somewhat puzzling. This apparent disparity motivated the current study. Using modern approaches and—importantly—by carefully controlling protein expression levels, our data directly contradict the selective retention model, and reveal that members from two distinct families of SV proteins are directly and specifically routed to axons.

The establishment of neuronal polarity and its maintenance has been studied for years, which prompts the question of why the direct and selective transport of SV proteins is only now being observed. As we show, without the careful control of expression levels, direct transport is obscured by the spillover and mistargeting of cargo into other compartments of neurons. We note that SV proteins seem to be particularly susceptible to this artifact, as the intentional overexpression of TfR did not appear to affect its selective transport to dendrites. In earlier studies, axonal proteins were likely delivered indiscriminately to axons and dendrites due to spillover and mistargeting, as a consequence of overexpression. Furthermore, there has been confusion in the field about what constitutes direct transport. Some groups defined direct transport according to where post-Golgi vesicles initially fuse (Nabb and Bentley 2022). By this definition, axonal cargo can leave the Golgi, traverse the entire neuron, including through dendrites, fuse in axons and still be considered a direct transport pathway. So, ultimately, this remains a version of the selective retention model.

Prior to the current study, there were reports in the literature that suggested a direct and selective transport pathway for SV proteins. Namely, an elegant study in nematode DA9 bipolar neurons demonstrated SYB2 was delivered directly to pre-synaptic boutons (P. Li et al. 2016). However, these neurons have a different microtubule organization than mammalian hippocampal neurons, so further studies were necessary to confirm whether this pathway extended to mammalian cells. Experiments in dorsal root ganglion (DRG) neurons suggested MAP2-dependent selective cargo sorting and transport of axonal proteins (Gumy et al. 2017), but these sensory neurons have pseudo-unipolar morphology and lacked an axon initial segment, thus making them distinct from hippocampal neurons. A more recent study, using mouse neurons, indicated that axonal proteins do not enter dendrites (Karasmanis et al. 2018). However, while dendritic exclusion was clearly established, entry into axons was not shown. Nevertheless, these papers began to question the idea of non-polarized transport.

New approaches have made it possible to directly address the trafficking of axonal cargos as they egress from the soma (Boncompain et al. 2012; Grimm et al. 2017; Los et al. 2008). At the outset of the current study, we showed that when expression levels are carefully controlled, two topologically distinct SV proteins egressed from the soma directly to axons without detouring through dendrites, thus uncovering a novel specific membrane transport pathway in neurons.

We used this controlled transport assay to conduct structure-function studies of SYT1, and found that glycosylation and palmitoylation were dispensable for direct transport. In contrast, removing the C2 domains of SYT1 disrupted polarized delivery to axons, and at steady state this mutant was present throughout the plasmalemma. We note that the HaloTag reporter was appended to the N-terminus of the truncation mutant but was on the C-terminus of the full-length protein and the PGM mutant. However, after careful design, tags were tolerated at each end of the full-length protein, so this is unlikely to affect localization (**Fig. 4, Fig. 5**).

This study also addressed the first half of the life cycle of SYT1 by conducting pulse-chase HaloTag assays to answer the long-standing question of whether SYT1 is—in fact—first delivered to the synaptic PM or pre-synaptic endosomes. Our results strongly argue that SYT1 is delivered to the presynaptic PM where it serves as a reservoir from which SVs are created. Then, SYT1 is internalized via its tandem C2-domains, potentially via mechanisms that mediate SV retrieval from the PM, during normal recycling. Retrieval may involve interactions with various adaptor proteins, but the emerging view is that this is unlikely to be occur at the PM, as clathrin-mediated endocytosis is no longer thought to mediate this step in recycling (Watanabe et al. 2013). Regardless, these pulse chase experiments reveal the first step in the biogenesis of SYT1-bearing SVs: selective delivery and incorporation of SYT1 in the presynaptic PM, as proposed decades ago (Buckley et al. 2000b; Feany et al. 1993; Hannah et al. 1999; Regnier-Vigouroux et al. 1991).

A key issue now is to understand the cargo selection process that underlies axon-specific transport, and to understand how newly delivered proteins are incorporated into SVs. Finally, a complete picture will not emerge until the other half of the life cycle of SV proteins is understood, namely how aged proteins are selected for, and undergo, degradation (Birdsall and Waites 2018; Cohen et al. 2013; Hoffmann-Conaway et al. 2020; Na et al. 2012; Sheehan et al. 2016; Sheehan and Waites 2017). New tools have made it possible to address these questions, including HaloTag pulse chase approaches, in conjunction with organelle isolation and mass spectrometry. These techniques promise to reveal, in biochemical detail, the itinerary of SV proteins as they are created and destroyed.

## Acknowledgements

We would like to thank the members of the Chapman lab, both past and present, for valuable discussion and feedback related to this manuscript. We thank K. Drerup for constructive comments regarding this study and M. Bradberry for the SYP-GFP fusion construct. This study was supported by grants from the NIH (MH061876 and NS097362 to E.R.C.). J.D.V. was supported by a postdoctoral fellowship from the National Institutes of Health F32 NS098604. E.R.C. is an Investigator of the Howard Hughes Medical Institute.

This article is subject to HHMI’s Open Access to Publications policy. HHMI lab heads have previously granted a nonexclusive CC BY 4.0 license to the public and a sublicensable license to HHMI in their research articles. Pursuant to those licenses, the author-accepted manuscript of this article can be made freely available under a CC BY 4.0 license immediately upon publication.

## Author Contributions

E.T.W.: conceptualization, data curation, formal analysis, investigation, methodology, project administration, supervision, validation, visualization, writing – original draft, writing – review and editing. M.M.P.: data curation, formal analysis, investigation, writing – review and editing. J.D.V.: conceptualization, methodology, supervision, writing – review and editing. E.R.C.: conceptualization, funding acquisition, methodology, project administration, resources, supervision, visualization, writing – original draft, writing – review and editing.

## Methods

### Ethics Statement

Animal care and use in this study were conducted in accordance with the NIH Guide for the Care and Use of Laboratory Animals handbook. Protocols were reviewed and approved by the Animal Care and Use Committee (ACUC) at the University of Wisconsin-Madison (Laboratory, Animal Welfare Public Health Service Assurance Number: A33688-01).

### Cell culture

Hippocampal neurons were dissected from pre-natal Sprague-Dawley rats on E18 (Envigo), or post-natal SYT1 conditional knockout floxed mice (Quadros et al. 2017) on P0-P1. Hippocampal tissue was maintained in chilled hibernate A media (BrainBits, HA) during dissection. After dissection, hippocampi were incubated in 0.25% trypsin (Corning, 25-053-CI) for 30 minutes at 37°C, triturated in DMEM (Thermo Fisher Scientific, 11965-118) supplemented with 10% fetal bovine serum (Atlanta Biological, S11550H) plus Penicillin-Streptomycin (Thermo Fisher Scientific, MT-30-001-CI), to dissociate tissue. Neurons were plated on poly-D-lysine (Thermo Fisher Scientific, ICN10269491) coated 18 mm coverslips (Warner instruments, 64-0734 (CS-18R17) at a density of 150,000 cells per coverslip in supplemented DMEM. Mouse hippocampal neurons were plated on 18 mm coverslips that were coated in poly-D-lysine and mouse laminin (Thermo Fisher Scientific, 23017015) for 2 hours at 37°C. Once the neurons settled (<1 hr), DMEM media was exchanged for Neurobasal-A (Thermo Fisher Scientific, 10888-022) medium supplemented with N21-MAX Media Supplement (R&D systems, AR008) (Chen et al. 2008), Glutamax (2 mM Gibco, 35050061), and Penicillin-Streptomycin. Additional supplemented Neurobasal-A media was added every 3-4 days to maintain the health of the cultures.

### Constructs

For the WT SYT1 (UniProt accession no. P21707) RUSH construct, a pre-prolactin leader sequence and streptavidin-binding peptide (SBP) were appended to its N-terminus, and a HaloTag (Promega, G7711) was fused to the C-terminus (**Fig. 1B**). Each of these moieties, in this and all other constructs, were attached via a flexible GS(GSS)_4_ linker. For the palmitoylation and glycosylation mutant form of SYT1, the palmitoylation sites, C74, C75, C77, C79, and C82 were substituted with Ala residues, and the glycosylation sites, T15/T16, and N24 were substituted with Ala and Gln residues, respectively. The truncated form of SYT1, SYT1ΔC2AB (a.a. 1-140), was generated in the same manner as the full-length protein, except that the HaloTag was placed at the N-terminus of the SYT1 coding sequence. For the SYB2 (UniProt accession no. P63045) RUSH construct, the HaloTag and SBP were appended to the C-terminus. For all SYT1 and SYB2 RUSH constructs, the HaloTag and the SBP were added in distinct positions to avoid steric interference between the SBP and the streptavidin hook. The streptavidin hook with an ER retention signal (Lys·Asp·Glu·Leu; KDEL) was made as a separate construct by sub-cloning from Str-KDEL_neomycin, a gift from F. Perez (Paris, France) (Addgene plasmid #65306; RRID:Addgene_65306) (Boncompain et al. 2012). A FLAG tag (DYKDDDDK) was added to the C-terminus of all RUSH constructs, immediately prior to the stop codon, to compare expression levels between co-expressed proteins. The TfR construct was kindly provided by J. Bonifacino (Bethesda, MD) (Chen et al. 2017). For the pulse-chase studies, a non-RUSH HaloTag-SYT1 construct was generated using the same pre-prolactin leader sequence as above, but now followed by an N-terminal HaloTag; for control experiments, the HaloTag was instead placed at the C-terminus. To mark synapses, a synaptophysin (SYP) GFP fusion protein (SYP-GFP), with the same flexible GS(GSS)_4_ linker between the two moieties, was used. All constructs were generated by overlap extension PCR and subcloned into the backbone using in-fusion cloning. Constructs were sequenced fully, and all maxi-preps were re-sequenced prior to use.

### Lentivirus production and use

Relevant constructs were subcloned into a pFUGW transfer plasmid (gift from D. Baltimore (Pasadena, CA); Addgene plasmid #14883; http://n2t.net/addgene:14883; RRID:Addgene_14883) (Lois et al. 2002). To make lentiviral expression neuron-specific, the ubiquitin promoter was replaced with a human synapsin I promoter (Kügler et al. 2003). Lentiviral particles were generated via calcium phosphate co-transfection of HEK293T cells (ATCC, CRL-3216; RRID:CVCL_0063) at 30-40% confluency with the pFUGW transfer plasmid and the packaging plasmids, pCD/NL-BH*DDD and pLTR-G. pCD/NL-BH*DDD was a gift from J. Reiser (Bethesda, MD) (Addgene plasmid #17531; http://n2t.net/addgene:17531; RRID:Addgene_17531) (Zhang et al. 2004), and pLTR-G was a gift from J. Reiser (Bethesda, MD) (Addgene plasmid #17532; http://n2t.net/addgene:17532; RRID:Addgene_17532) (Reiser et al. 1996). HEK293T cells were maintained in DMEM supplemented with 10% FBS and Penicilin-Streptomycin. Following transfection, the supernatant was collected after 48 hours of expression, filtered with a 0.45 mm PVDF filter to remove cells and large cell debris, and concentrated by ultracentrifugation at 110,000 x g for two hours. Viral particles were re-suspended in Ca^2+^/Mg^2+^-free phosphate buffered saline (PBS), aliquoted, and stored at -80°C (Kutner et al. 2009).

For RUSH release experiments, neurons were transduced with the streptavidin hook virus on 8 DIV and transduced with a reporter virus on 9 DIV. In Fig. 1 and Fig. 2, a virus that expressed GFP with a “KDEL” retention-signal on the C terminus to label ER, was also transduced on 9 DIV. Cells were imaged on 14-16 DIV. Lentivirus was titrated based on fluorescence and coverage unless otherwise stated in the text.

### Transfection

For pulse-chase experiments, neurons were cultured in 12-well plate cell culture plates (Genesee Scientific; 25-106) and co-transfected with SYP-GFP and SYT1-HaloTag or HaloTag-SYT1 on 9 DIV using Lipofectamine LTX Reagent with PLUS Reagent (Thermo Fisher Scientific, 15338-100). Briefly, DNA plasmids were diluted in 25 μl Opti-MEM I Reduced Serum Medium (Gibco; 31985062), then 0.25 μl PLUS reagent was added. Separately, 1 μl LTX Reagent was diluted in 25 μl of Opti-MEM I. The DNA-PLUS reagent mixture was added dropwise to the LTX reagent mixture then added to culture media in each well.

### Janelia Fluor dye usage

HTL-conjugated JF dyes were graciously provided by L. Lavis (Ashburn, VA). We made use of JF549, and JF549i. For protein localization of the RUSH constructs, cultures were incubated with 100 nM JF549 for 30-60 minutes at 37°C then rinsed twice prior to imaging. For concurrent ICC experiments, the JF dye JF549 was added to the secondary antibody mix and incubated at 25°C for 1 hour. For the live-cell pulse-chase labeling experiments, cultures were incubated with 1 nM JF549i for 6 days at 37°C, rinsed twice, and imaged in an environmental chamber. Incubation with JF549i for up to 8 days showed no detectable nonspecific uptake of this dye or crossing of the PM. JF549 was added to the coverslip at a final concentration of 100 nM during imaging.

### Live-cell imaging

Prior to imaging, RUSH reporter proteins were labeled with JF549 HTL (Janelia Farms) and, for rat neurons, anti-pan-neurofascin antibody (UC Davis/NIH NeuroMab Facility, A12/18; RRID:AB_2877334) for 60 minutes. Coverslips were rinsed twice with warmed PBS and returned to conditioned NBM growth media. Coverslips were incubated with IgG2α Alexa Fluor 647 secondary (Thermo Fisher Scientific, A-21241; RRID:AB_2535810) for 15-30 minutes to label the anti-pan-neurofascin primary antibody. Coverslips were rinsed twice with warmed PBS and imaged in standard ECF imaging solution (140 mM NaCl, 5 mM KCl, 2 mM CaCl_2_, 2 mM MgCl_2_, 5.5 mM glucose, 20 mM HEPES (pH 7.3) in PBS) at 37°C. The reporter proteins were released from the ER-localized streptavidin ‘hook’ with the addition of 40 μM biotin (Sigma-Aldrich, B4639-100MG) to the coverslip. Biotin was diluted in 200 μl of ECF imaging solution and added to 800 μl of media in the imaging chamber for a final concentration of 40 μM. Videos were acquired ∼20-30 minutes after biotin addition at 1 frame per second with the Zeiss 880 Airyscan LSM microscope and 63x objective using Fast Airyscan mode. All images were processed with automatic Airyscan deconvolution settings. Temperature, CO_2_, and humidity were controlled using an Oko-lab incubation system.

### Kymograph generation and analysis

Kymographs (20 μm) were generated from 60 second RUSH movies from the soma-out direction using ZEN blue software 3.0 (ZEISS; Oberkochen, Germany), and were analyzed manually in Fiji (Schindelin et al. 2012). Directionality, as well and distance and time parameters, were recorded for each vesicle movement identified in the kymographs. For all figures, kymograph lines with a negative slope represent anterograde transport, and those with a positive slope indicate retrograde transport. The kymographs analyzed in Fig. 4 were blinded prior to analysis.

### Immunocytochemistry (ICC)

Dissociated cultures were fixed with 4% paraformaldehyde, permeabilized with 0.2% saponin, and then blocked (0.04% saponin, 10% goat serum and 1% BSA in PBS), and immunostained at 4°C (0.1% BSA and 0.04% saponin in PBS) overnight. The following morning coverslips were rinsed 3 times for 5-minute intervals with PBS and incubated with secondary antibodies (0.1% BSA and .04% saponin in PBS) for 1 hour. Coverslips were rinsed 3 times for 5 minute intervals with PBS and mounted on microscope slides (Fisher Scientific, 22-178277) using ProLong Glass Antifade Mountant (Thermo Fisher Scientific, P36980) or ProLong Glass Antifade with Mountant with NucBlue Stain (Thermo Fisher Scientific, P36981).

### Protein Immunoblots (IB)

Neuronal cell lysates were collected from dissociated neuronal cultures with 150 μl lysis buffer (2% SDS, 1% Triton X-100, and 10mM EDTA in PBS, plus (1:200) 250 mM PMSF, and (1:500) 1 mg/ml aprotinin, leupeptin, and pepstatin A protease inhibitors. Samples were boiled at 100°C for 5 minutes after the addition of 50 μl of sample buffer (DTT, glycerol, and bromophenol blue) and 20 μl of lysates were run on 13.5% acrylamide gels with 10% 2,2,2,-Trichloroethanol (TCE) (Sigma-Aldrich; T54801-100G). After protein separation by SDS-PAGE, the TCE was activated by UV light (300 nm) and the cross-linked proteins were imaged with the ChemiDoc MP Imaging System (Bio-Rad Laboratories) as a loading control (Ladner et al. 2004). SDS-PAGE gels were transferred to a PVDF membrane (Immobilon-FL; EMD Millipore) for 30 minutes per gel at a constant 240 mA, then blocked with 5% nonfat milk protein in Tris-buffered saline plus 1% Tween 20 (TBST) for 30 minutes. PVDF membranes were incubated in primary antibody and 1% milk in TBST overnight at 4°C. The next day the membrane was rinsed and incubated with a secondary antibody in 1% milk in TBST for 1 hour then washed three times for a total of 15 minutes. All washes were done with TBST. Immunoblots were imaged using Luminata Forte Western HRP substrate (EMD Millipore; ELLUF0100) and the ChemiDoc MP Imaging System (Bio-Rad Laboratories). Bands were analyzed by densitometry and contrast was linearly adjusted for publication using Fiji (Schindelin et al. 2012).

### Statistics

Exact values from experiments and analysis, including the number of data points (n) and number of trials are included in the figure or are listed in the figure legends. Analysis was performed using GraphPad Prism 9.20 (GraphPad Software Inc.). Normality was assessed by histograms of data and QQ plots; if normal, parametric statistical methods were used, if not, nonparametric methods were used for analysis.

### Antibodies

Primary Antibodies
| Antibody | Source | Identifier | Concentration |
| --- | --- | --- | --- |
| anti-SYT1 (48) | Developmental Studies Hybridoma Bank | Cat# mAB 48 (asv 48)<br>RRID:AB_2199314 | ICC (1:500)<br>IB (1:500) |
| anti-SYB2/VAMP (69.1) | Synaptic Systems | Cat# 104211<br>RRID:AB_2619758 | ICC (1:500)<br>IB (1:1K) |
| anti-GM130 | BD Transduction Laboratories | Cat# 610822<br>RRID AB_398142 | ICC (1:500) |
| anti-pan-neurofascin (extracellular) antibody (A112/18) | NeuroMab | Cat# 75-172<br>RRID:AB_2282826 | ICC (1:1K) |
| anti-SYP | Cedarlane Labs | Cat# 101004(SY)<br>RRID:AB_1210382 | ICC (1:500)<br>IB (1:1K) |
| anti-MAP2 | Sigma-Aldrich | Cat# AB5543<br>RRID:AB_571049 | ICC (1:250) |

Secondary Antibodies
| Antibody | Source | Identifier | Concentration |
| --- | --- | --- | --- |
| goat anti-mouse<br>IgG2α-Alexa Fluor 647 | Thermo Fisher<br>Scientific | Cat# A21241<br>RRID:AB_2535810 | 1:500 |
| goat anti-mouse<br>IgG2β-Alexa Fluor 647 | Thermo Fisher<br>Scientific | Cat# A21242<br>RRID:AB_2535810 | 1:500 |
| goat anti-guinea pig<br>IgG-Alexa Fluor 647 | Thermo Fisher<br>Scientific | Cat# A21450<br>RRID:AB_2735091 | 1:500 |
| goat anti-chicken<br>IgG-Alexa Fluor 405 | Abcam | Cat# ab175675<br>RRID:AB_2810980 | 1:500 |
| goat anti-mouse<br>IgG-HRP | Bio-Rad Laboratories | Cat# 1706516<br>RRID:AB_11125547 | 1:10K |
| goat anti-mouse<br>IgG2b-HRP | Bio-Rad Laboratories | Cat# M32407<br>RRID:AB_2536647 | 1:10K |
| goat anti-guinea pig<br>IgG-HRP | Abcam | Cat# ab6908<br>RRID:AB_955425 | 1:10K |

## Resource Availability

### Lead Contact

Further information and requests for resources and reagents should be directed to and will be fulfilled by the Lead Contact, Dr. Edwin Chapman.

### Materials Availability

All unique/stable reagents generated in this study are available from the Lead Contact with a completed Materials Transfer Agreement.

## Supplementary Figures

**Figure S1.**
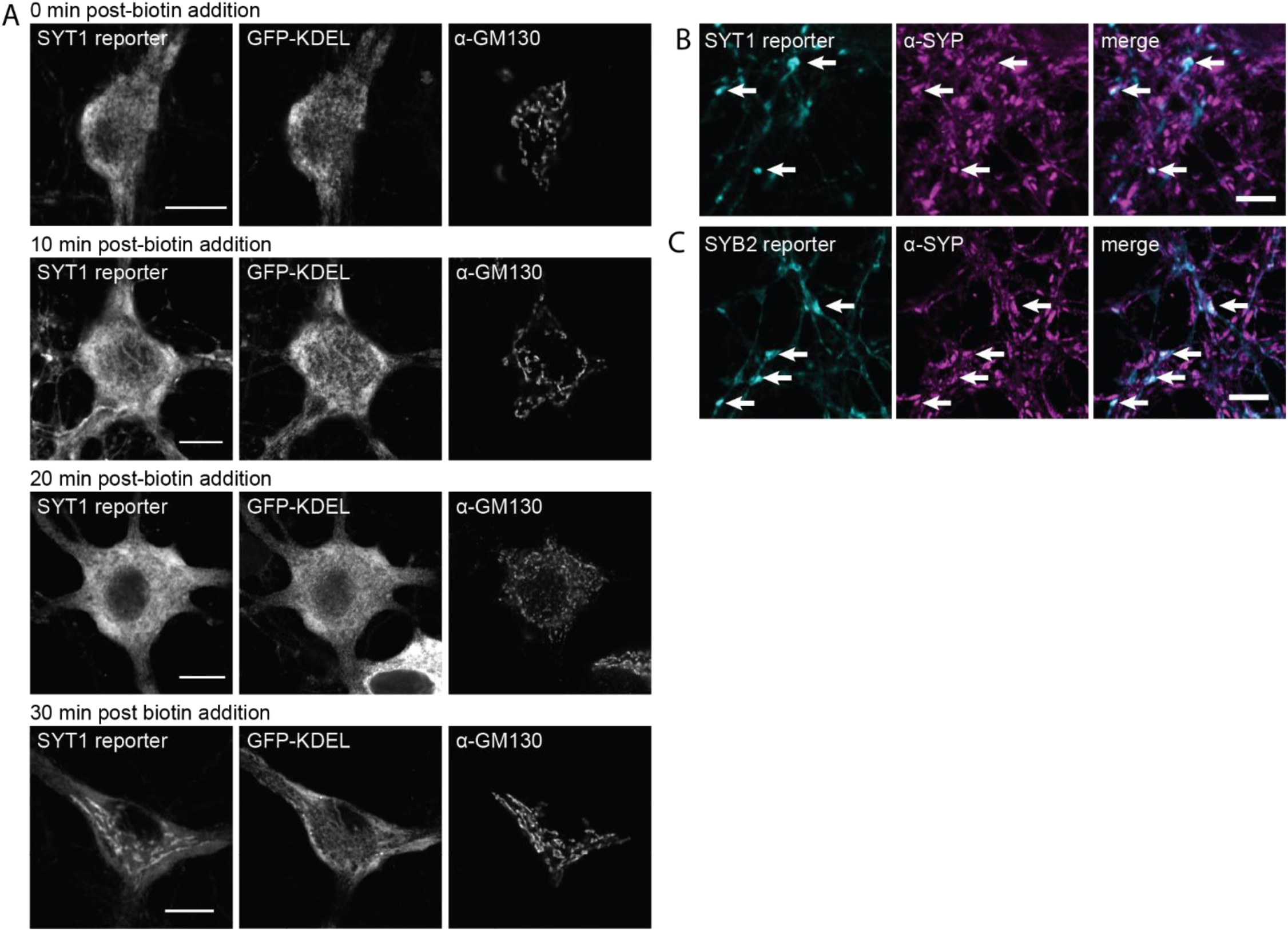
The RUSH system (**A**) Super-resolution, fixed-cell optical sections of 15 DIV rat hippocampal neurons expressing the SYT1 reporter and ER-targeted GFP, detailing the movement of the SYT1 reporter from the ER to the Golgi after biotin addition (0, 10, 20, and 30 minutes), as indicated. Scale bars represent 10 μm. (**B, C**) Same as panel (**A**), but stained for the SYT1 reporter and SYB2 reporter, respectively, and α-synaptophysin (α-SYP) to confirm proper targeting to synapses. Note that all neurons were stained for SYP, but only a handful of cells expressed the SYT1 or SYB2 reporter. Arrows denote colocalization. Scale bars represent 5 μm.

**Figure S2.**
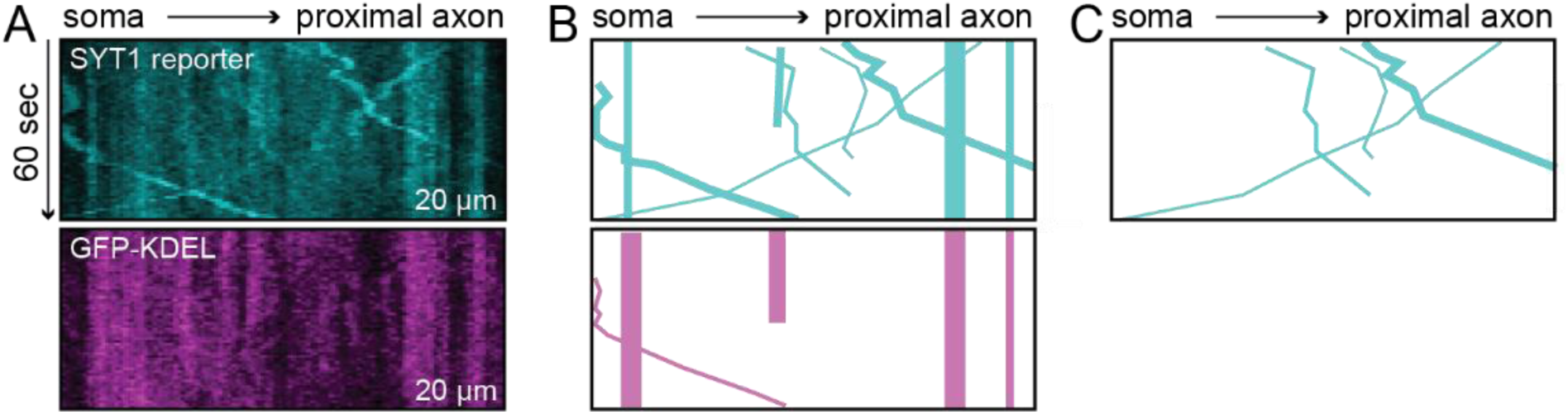
Kymograph analysis (**A**) A representative kymograph of the SYT1 reporter in proximal axons. The SYT1 channel, visualized by HaloTag and the JF549 ligand, and the ER-targeted GFP (GFP-KDEL) channel, are shown. (**B**) Vesicles identified in the kymographs were traced in both channels. We note that there is significant labeling of the ER in the GFP-KDEL kymograph and thus traced only the areas that overlapped with the SYT1 reporter for the sake of clarity. (**C**) The movement of vesicles harboring the SYT1 reporter were analyzed; vesicles that also carried GFP-KDEL were excluded.

**Figure S3.**
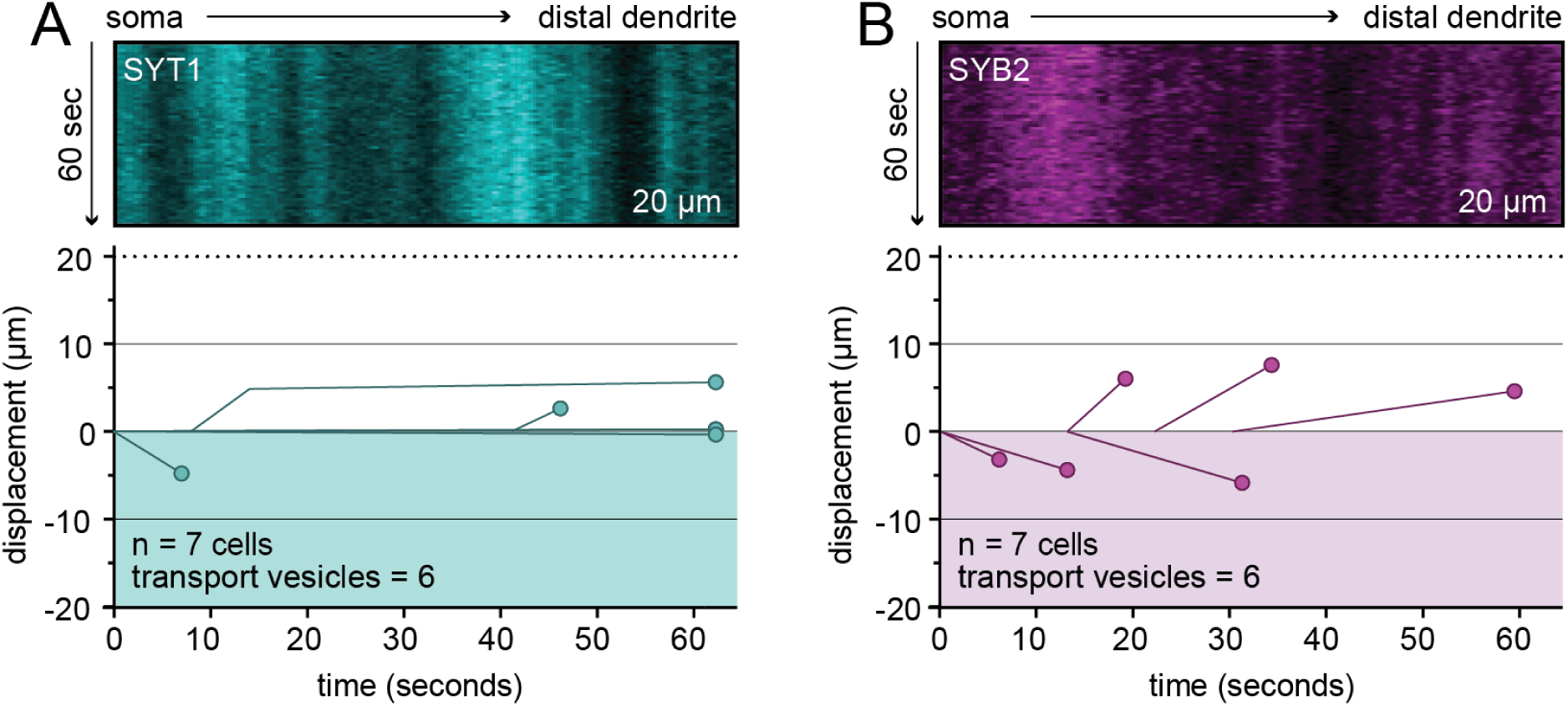
SV RUSH constructs in distal dendrites Representative kymographs from the distal dendrite of rat hippocampal cultures at 14-16 DIV expressing the SYT1 (**A**) or SYB2 (**B**) reporters. Few vesicles were observed in distal dendrites, despite the fact that the total number of cells, time, and distance observed was the same as for axons, where robust activity was detected.

**Figure S4.**
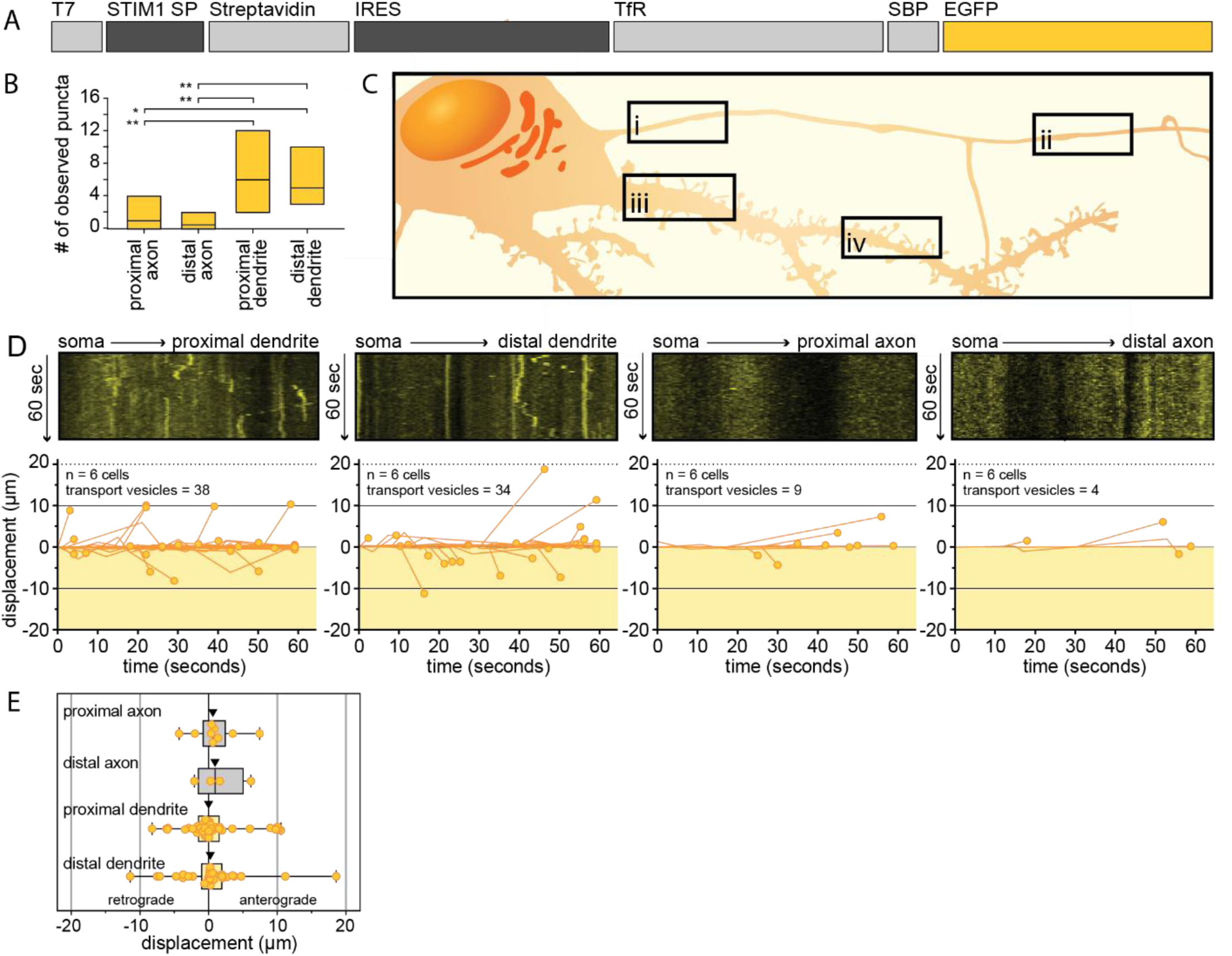
RUSH of TfR reporter in rat hippocampal neurons (**A**) A schematic of the bicistronic pIRES vector used to express the TfR reporter and streptavidin hook protein in the same cell. (**B**) The number of TfR-bearing transport vesicles observed in a cell for 1 minute in each region was plotted as mean values <u>±</u> SD for proximal axons (1.50 ± 1.6 puncta), distal axons (0.667 ± 0.82 puncta), proximal dendrites (6.33 ± 3.5 puncta), and distal dendrites (5.67 ± 2.3 puncta). Data were collected from 6 cells from 5 litters. A one-way ANOVA with multiple comparisons was run, and p-values were as follows: proximal axon vs. distal axon = 0.79; proximal axon vs. proximal dendrite = 0.0064; proximal axon vs. distal dendrite = 0.015; distal axon vs. proximal dendrite = 0.0022; distal axon vs. distal dendrite = 0.0060; proximal dendrite vs. distal dendrite = 0.79. (**C**) An illustration outlining proximal axons (i), distal axons (ii), proximal dendrites (iii), and distal dendrites (iv) that were imaged for analysis. These proximal and distal regions follow the same definitions described for **Fig. 2**. All kymographs were 20 μm in length. (**D**) Representative kymographs from rat hippocampal neurons (14-16 DIV), after release of the tethered TfR reporter, for each compartment are shown, with the data quantified and plotted immediately below. The number of cells and transport vesicles are also indicated. Kymographs, and the corresponding displacement graphs, from proximal and distal dendrites demonstrate robust transport of the TfR reporter; in contrast, axons lacked TfR transport. (**E**) Displacement of transport vesicles containing the TfR reporter were plotted in the anterograde (positive) or retrograde (negative) direction with respect to the soma for each neurite (arrowheads indicate median; proximal axon = 0.889 μm, distal axon = 1.47 μm, proximal dendrite = 0.558 μm, distal dendrite = 0.318 μm).

**Figure S5.**
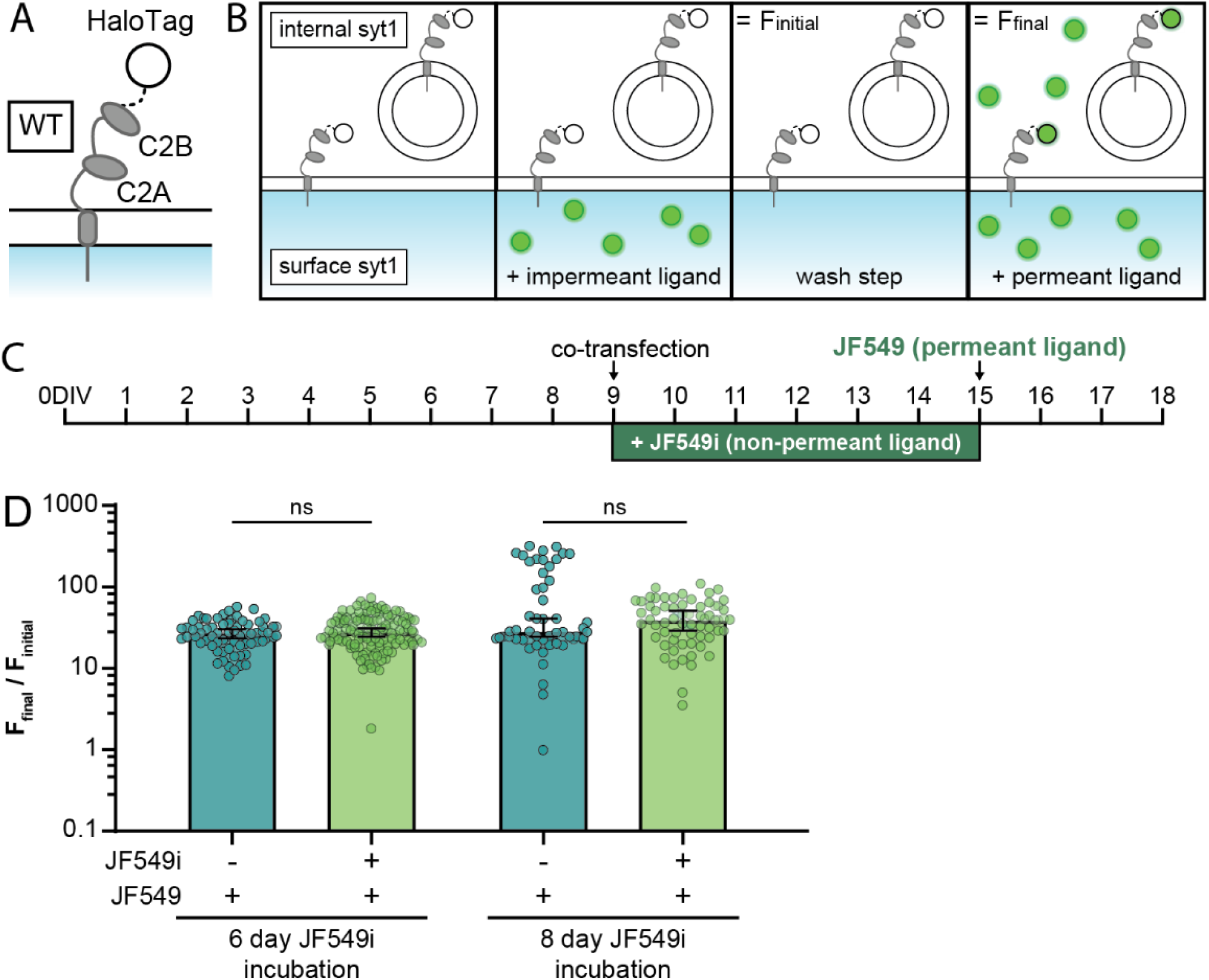
Testing the permeability of JF549i (**A**) A cartoon of SYT1 with a C-terminal HaloTag and (**B**) the time course of ligand addition during the pulse-chase assay. By appending the HaloTag to the cytoplasmic domain, the tag is not exposed to the extracellular milieu, and should not be labeled with non-permeant JF549i ligand. (**C**) Timeline for transfecting and labeling neurons. This scheme is the same as the experiment conducted in **Fig. 5**, but—again—with the HaloTag oriented inside the cell when SYT1 is in the PM. (**D**) Plots of the change in fluorescence (F_final_/F_initial_) upon adding a permeant fluorescent ligand relative to cells grown with or without non-permeant ligand for 6 or 8 days. Median values, with 95% CI, are shown. These values were: 6 days with (27.87, [24.37, 30.79]) and without JF549i (27.11, [23.42, 30.37]), or 8 days with (39.40, [28.87, 50.64]) and without JF549i (28.27, [24.41, 40.95]). A Mann-Whitney test was run for both 6- (p=0.260) and 8-day (p=0.890) incubation conditions. No difference in F_final_/F_initial_ between cultures grown with and without the non-permeant ligand, was observed. Thus, incubation with the JF549i ligand did not result in any significant labeling of the cytoplasmic HaloTag and is non-permeant under these experimental conditions. Data were collected as follows, with synapse, fields of view, and number of litters listed in order: 6-day incubation with JF549i: 125, 3, 1; 6-day incubation without JF549i: 68, 4, 1; 8-day incubation with JF549i: 60, 3, 1; 8-day incubation without JF549i: 51, 3, 1.

**Table S1.** Descriptive statistics corresponding to Fig. 2. (**A**) Corresponds to **Fig. 2H** and **Fig. 2I**. Table (**B**) correspond to **Fig. 2J** and **Fig. 2K**.

|  | SYT1 reporter |  |  |  |  | SYB2 reporter |  |  |  |
| --- | --- | --- | --- | --- | --- | --- | --- | --- | --- |
|  | Proximal axon | Distal axon | Proximal dendrite | Distal dendrite |  | Proximal axon | Distal axon | Proximal dendrite | Distal dendrite |
| Number of values (transport vesicles) | 29 | 21 | 4 | 6 |  | 26 | 27 | 10 | 6 |
| Mean | 6.59 | 10.1 | -1.68 | 1.16 |  | 5.33 | 4.27 | -0.0789 | 2.27 |
| Median | 8.69 | 11.7 | -0.179 | 0.255 |  | 5.07 | 4.60 | 0.438 | 4.47 |
| Std. Deviation | 9.1 | 8.4 | 3.8 | 3.0 |  | 8.3 | 8.7 | 8.0 | 5.4 |
| Std. Error of Mean | 1.7 | 1.8 | 1.9 | 1.4 |  | 1.6 | 1.7 | 2.5 | 2.2 |
| Lower 95% CI of mean | 3.11 | 6.21 | -7.75 | -2.65 |  | 2.00 | 0.839 | -5.81 | -3.41 |
| Upper 95% CI of mean | 10.1 | 13.9 | 4.40 | 4.97 |  | 8.66 | 7.70 | 5.66 | 7.95 |

**Table S1 - Descriptive statistics corresponding to Fig. 2**
|  | SYT1 reporter |  |  |  |  | SYB2 reporter |  |  |  |
| --- | --- | --- | --- | --- | --- | --- | --- | --- | --- |
|  | Proximal axon | Distal axon | Proximal dendrite | Distal dendrite |  | Proximal axon | Distal axon | Proximal dendrite | Distal dendrite |
| Number of values (transport vesicles) | 7 | 7 | 7 | 7 |  | 7 | 7 | 7 | 7 |
| Mean | 4.14 | 3.00 | 0.571 | 0.857 |  | 3.71 | 3.86 | 1.43 | 0.857 |
| Median | 4.0 | 4.0 | 1.0 | 1.0 |  | 5.0 | 4.0 | 2.0 | 1.0 |
| Std. Deviation | 2.0 | 2.4 | 0.53 | 0.69 |  | 2.0 | 2.2 | 0.98 | 0.69 |
| Std. Error of Mean | 0.77 | 0.90 | 0.20 | 0.26 |  | 0.75 | 0.83 | 0.37 | 0.26 |
| Lower 95% CI of mean | 2.26 | 0.80 | 0.077 | 0.22 |  | 1.89 | 1.83 | 0.526 | 0.219 |
| Upper 95% CI of mean | 6.03 | 5.20 | 1.07 | 1.50 |  | 5.54 | 5.89 | 2.33 | 1.50 |

**Table S2.** Descriptive statistics corresponding to Fig. 4. (**A**) Corresponds to **Fig. 4I**. (**B**) A breakdown of transport vesicle movements (fraction of a whole) related to **Fig. 4J**.

| | SYT1 | | SYT1ΔC2AB | | SYT1-PGM | |
| --- | --- | --- | --- | --- | --- | --- |
|  | Axon | Dendrite | Axon | Dendrite | Axon | Dendrite |
| Number of values (transport vesicles) | 10 | 10 | 11 | 11 | 8 | 8 |
| Mean | 6.60 | 2.10 | 5.91 | 3.64 | 6.25 | 1.88 |
| Median | 7.5 | 1.5 | 6.0 | 3.0 | 5.5 | 1.5 |
| Std. Deviation | 4.2 | 1.8 | 3.1 | 2.3 | 3.4 | 1.8 |
| Std. Error of Mean | 1.3 | 0.57 | 0.94 | 0.70 | 1.2 | 0.64 |
| Lower 95% CI of mean | 3.62 | 0.818 | 3.82 | 2.07 | 3.43 | 0.364 |
| Upper 95% CI of mean | 9.58 | 3.38 | 8.00 | 5.21 | 9.067 | 3.39 |

**Table S2 - Descriptive statistics corresponding to Fig. 4**
| | SYT1<sub>axon</sub> | SYT1<sub>dendrites</sub> | ΔC2AB<sub>axon</sub> | ΔC2AB<sub>dendrites</sub> | PGM<sub>axon</sub> | PGM<sub>dendrite</sub> |
| --- | --- | --- | --- | --- | --- | --- |
| anterograde | 0.517 | 0.259 | 0.433 | 0.261 | 0.403 | 0.633 |
| retrograde | 0.117 | 0.444 | 0.192 | 0.329 | 0.229 | 0.167 |
| retrograde with pause/reverse | 0.019 | 0.000 | 0.000 | 0.013 | 0.041 | 0.000 |
| anterograde with pause/reverse | 0.041 | 0.019 | 0.071 | 0.091 | 0.021 | 0.083 |
| stationary (1<5μm) | 0.067 | 0.028 | 0.077 | 0.024 | 0.000 | 0.033 |
| stationary (<1μm) | 0.240 | 0.250 | 0.227 | 0.281 | 0.306 | 0.083 |

**Table S3.** Descriptive statistics corresponding to Fig. 5. (**A**) Corresponds to **Fig. 5E**. (**B**) Corresponds to **Fig. 5F**.

|  | Cultured in the absence of JF549i |  | Cultured in the presence of JF549i |  |
| --- | --- | --- | --- | --- |
|  | Before permeant ligand addition | After permeant ligand addition | Before permeant ligand addition | After permeant ligand addition |
| Number of values (synapses) | 136 | 136 | 156 | 156 |
| Mean | 54.2 | 1222 | 1389 | 2563 |
| Median | 41.5 | 1115 | 1170 | 2140 |
| Std. Deviation | 79.1 | 624.3 | 1274 | 2080 |
| Std. Error of Mean | 6.78 | 53.5 | 102.0 | 166.7 |
| Lower 95% CI of mean | 40.8 | 1116 | 1188 | 2234 |
| Upper 95% CI of mean | 67.6 | 1328 | 1591 | 2892 |

**Table S3 - Descriptive statistics corresponding to Fig. 5**
|  | Cultured in the absence of JF549i | Cultured in the presence of JF549i |
| --- | --- | --- |
| Number of values (synapses) | 136 | 156 |
| Mean | 30.5 | 2.03 |
| Median | 25.4 | 1.82 |
| Std. Deviation | 21.6 | 0.925 |
| Std. Error of Mean | 1.85 | .0740 |
| Lower 95% CI of mean | 26.8 | 1.88 |
| Upper 95% CI of mean | 34.2 | 2.17 |

